# Genome-Wide Prediction of *cis*-Regulatory Regions Using Supervised Deep Learning Methods

**DOI:** 10.1101/041616

**Authors:** Yifeng Li, Wenqiang Shi, Wyeth W. Wasserman

## Abstract

Identifying active *cis*-regulatory regions in the human genome is critical for understanding gene regulation and assessing the impact of genetic variation on phenotype. Based on rich data resources such as the Encyclopedia of DNA Elements (ENCODE) and the Functional Annotation of the Mammalian Genome (FANTOM) projects, we introduce DECRES, the first supervised deep learning approach for the identification of enhancer and promoter regions in the human genome. Due to their ability to discover patterns in large and complex data, the introduction of deep learning methods enables a significant advance in our knowledge of the genomic locations of *cis*-regulatory regions. Using models for well-characterized cell lines, we identify key experimental features that contribute to the predictive performance. Applying DECRES, we delineate locations of 300,000 candidate enhancers genome wide (6.8% of the genome, of which 40,000 are supported by bidirectional transcription data) and 26,000 candidate promoters (0.6% of the genome).

---

In this article, we apply deep supervised analysis methods to identify the positions of active *cis*-regulatory regions (CRRs), including both enhancers and promoters, across the human genome. CRRs play a crucial role in precise control of gene expression. Promoters and enhancers act via complex interactions across time and space in the nucleus to control when, where and at what magnitude genes are active. CRRs, through interactions with proteins such as histones and sequence-specific DNA-binding transcription factors (TFs), help specify the formation of diverse cell types and respond to changing physiological conditions. While gene expression is ultimately a reflection of regulation across multiple processes, the key role of promoters and enhancers has been a central focus of genome annotation for the past decade. The investment in generating informative data for the detection of these regions has been immense, in part motivated by the anticipation that advanced computational approaches would be able to transform the data into a reliable annotation of the genome.

Promoters and enhancers were early discoveries during the molecular characterization of genes. While promoters specify and enable the positioning of RNA polymerase machinery at transcription initiation sites, enhancers modulate the activity of promoters from linearly distal locations away from transcript initiation sites [1, 2]. The delineation between the classes has become increasingly challenging, with some literature suggesting the two categories are the edges of a continuous spectrum of CRRs [3]. Indeed, it has long been observed that sequences flanking transcription initiation regions can function as enhancers (promoter-proximal regions), and in recent years, it has been observed that there are transcripts initiated at the edges of active enhancers [4, 5]. For the purpose of this report, we address the two as distinct classes, but discuss the relationship between our findings and the continuous class model.

The use of computational methods to detect the locations of promoters and enhancers has been a key focus of bioinformatics for twenty years (see reviews [6] and [7]). With the advances of experimental procedures for profiling the properties of chromatin and RNA transcripts, a new wave of methods has arrived. Given the small set of reliable enhancer annotations, it was appropriate that the first among these methods used unsupervised learning. For instance, both ChromHMM [8] and Segway [9] segment the genome into sequence classes based on ENCODE project data [10], such as histone modification ChIP-seq (chromatin immunoprecipitation followed by sequencing [11]) signals. Such unsupervised methods infer hidden states based on observed signals, and then associate an element to each hidden state. The states are subsequently labelled with biological functions based on enrichment for known examples. A test of predicted Enhancers for the K562 leukemia cell line by the Combined method (unifying ChromHMM and Seqway annotations) [12] using a high-throughput reporter gene assay [13] revealed that only 26% of predicted enhancers have regulatory activity [14]. The assessment showed that the predicted Weak Enhancers, a class associated with lower H3K27ac and H3K36me3 signals, unexpectedly drove higher gene expression than the predicted Enhancers. It is evident that improvements are needed, potentially involving the use of additional experimental features and alternative machine learning approaches.

Despite the limited set of precisely annotated active enhancers, supervised machine learning models have been attempted to predict enhancer regions. In each case, a distinct definition of a suitable positive training set of enhancers was taken. A random-forest method was used in [15] to classify TF bound regions with a focus on observed binding patterns, generating sets of two-class classifiers to distinguish regions based on binding activity and position relative to promoter regions. A random-forest based enhancer classification method was devised in [16] with histone modification ChIP-seq data as features, using p300 bound regions as the basis for training. Chen *et al.* applied multinomial logistic regression with LASSO regularization to find key features for the classification of stem cell-specific functional enhancer regions [17]. Using STARR-seq data, a new experimental approach for screening candidate enhancer sequences [18], dinucleotide repeat motifs (DRMs) were found to be enriched in broadly active enhancers, leading to a proposition that a small set of TF binding site motifs and DRMs might be sufficient for enhancer prediction [19].

New laboratory methods are emerging, providing a refined resolution of CRR locations. The majority of human DNA is transcribed, producing diverse types of RNA. In particular, transcripts generated at the edges of enhancers, enhancer RNAs (eRNAs), allow for the experimental readout of active regulatory regions. Global run-on and sequencing (GRO-seq) protocols [20] measure the 5’-end of nascent RNAs revealing the divergent transcriptional signature of both transcriptionally active promoters and enhancers [5]. Using GRO-seq signals, a support vector regression model (dReg) was developed to predict active transcriptional regulatory elements [21]. The cap analysis of gene expression (CAGE) technique [22] captures the 5’-end of RNA transcripts, enabling a precise determination of transcript initiation sites. Using CAGE, the FANTOM5 Consortium has identified an atlas of transcriptionally active promoters [23] and a permissive set of 43,011 transcriptionally active enhancers characterized by bidirectional eRNAs [4] across hundreds of human cell types and tissues. These enhancers were validated with high success rates ranging from 67.4% to 73.9% [4]. Compared to protein-coding RNAs, eRNAs are believed to degenerate quickly, and only a small number of tissues have been explored with sufficient depth to reveal eRNAs. While the FANTOM enhancer set is therefore incomplete, it provides a uniquely large inventory of high-quality enhancers to use for the training of machine learning approaches. An ensemble support vector machine method suggested the potential to distinguish enhancers based on such data [24].

We have previously proposed and herein present the use of a deep feature selection (DFS) model for the supervised prediction of CRRs [25]. Deep learning is a dramatic advance in the frontier of artificial intelligence [26, 27, 28]. Unlike widely used linear models, deep learning approaches model complex systems and capture human-understandable patterns. Driven by big and rich data, deep learning has been successfully applied in various areas such as automatic image annotation and speech language processing [29]. Bioinformaticians have started using this powerful tool for next-generation sequencing data mining, such as predicting the impact of variations on exon splicing [30], detecting TF binding patterns [31], and predicting protein secondary structures [32].

Our study stands on three important legs. First, the precisely annotated FANTOM promoters and enhancers, which provide the largest experimentally defined collection of CRRs. Second, the ENCODE project genome-wide feature data, such as histone modifications, TF binding, RNA transcripts, chromatin accessibility, and chromatin interactions. Third, deep learning methods to distinguish CRRs based on the available data. We unite the three components to create the DECRES model, with which we identify the most comprehensive collection of CRRs across the human genome yet compiled.

## Results

### Deep learning accurately distinguishes active enhancers and promoters from background

We investigated the capacity of deep learning models to separate enhancers and promoters, and to distinguish them from other regions and between activity states. Training a deep three-layer feedforward neural network over our labelled data from well-characterized cells (see Methods, Supplementary Tables 1 and 2), we recorded the mean sensitivity, specificity, and overall accuracy (using 10-fold cross-validation) in Figure 1 and Supplementary Figure 1. For narrative convenience, hereafter we refer to active enhancer, active promoter, active exon, inactive enhancer, inactive promoter, inactive exon, and unknown (or uncharacterized) region as A-E, A-P, A-X, I-E, I-P, I-X, and UK, respectively. Under the assumption that active CRRs are undergoing transcription, active applies to regions in which CAGE transcript initiation events are observed in the tissue of focus, while inactive refers to regions detected in other tissues, but not in the focus tissue. First, we are able to distinguish between active enhancers and promoters (A-E versus A-P) (Fig. 1A). We used A-E and A-P as positive and negative training classes, respectively. Overall, we found that A-E and A-P are highly separable, with the highest mean accuracy of 93.59% on GM12878 lymphoblastoid cells, and the lowest of 87.78% and 88.14% on MCF7 and A549 cells, respectively (results correlate with smaller training sample sizes). Second, we can distinguish active and inactive CRRs (either enhancers or promoters). From Figure 1B and Supplementary Figure 1A, it can be observed that accuracies on GM12878, HelaS3, HepG3, and K562, which have the largest training sets, are above 90% with small variances for both enhancers and promoters. Consistent with larger promoter training data, all sets exceeded 90% accuracy, while for the smaller enhancer sets, from A549 and MCF7 cells, lower accuracies with larger variances were obtained. In the rest of this paper, we exclude A549 and MCF7 cell lines due to limited data availability. Third, not unexpectedly, it is difficult to distinguish between inactive enhancers and promoters (Supplementary Fig. 1B). None of the mean accuracies for the eight cell types exceeded 80%. As the properties of inactive sequences could be similar for both enhancers and promoters in a cell of focus, we elected to group the inactive CRRs and assess the capacity of our deep method to distinguish them from active CRRs. When grouping CRRs and seeking to distinguish activity states, we obtained a mean accuracy higher than 90% across cells, for example 95.87% on GM12878. It is evident that our deep approach has a strong capacity to distinguish between classes when there is both sufficient training data and when the underlying biological classification is appropriate. Fourth, we tested the applicability of predicting A-E and A-P from the super background (BG) class merging I-E, I-P, A-X, I-X, and UK (Fig. 1C). The results are promising. If A-E and A-P are merged further to form a super class (A-E+A-P), higher performance is achieved (Supplementary Fig. 1C).

**Figure 1:**
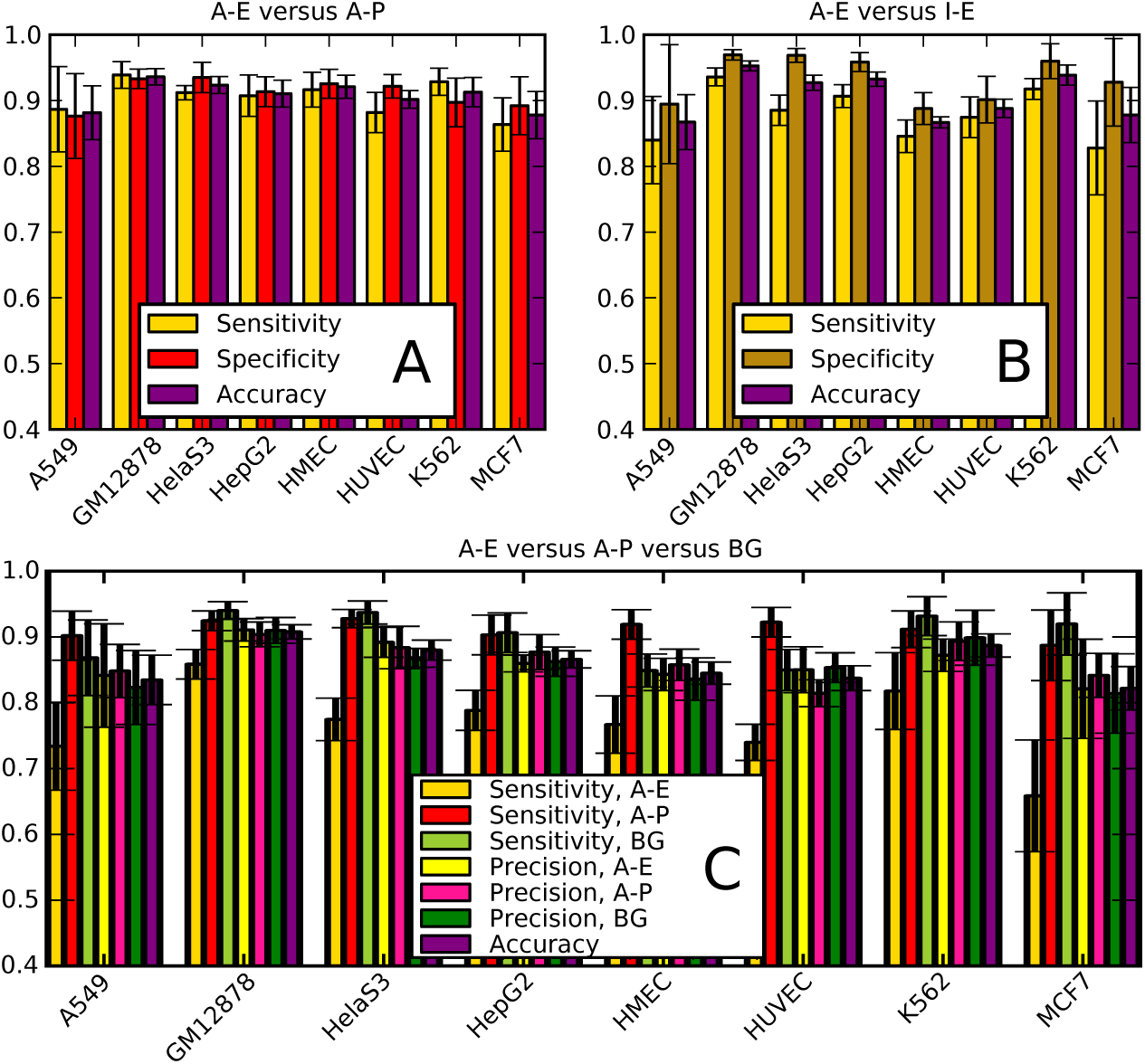
The mean performance and standard deviation of 10-fold cross-validations using the MLP model on our labelled data of eight cell types. A-E: Active Enhancer, A-P: Active Promoter, A-X: Active Exon, I-E: Inactive Enhancer, I-P: Inactive Promoter, I-X: Inactive Exon, UK: Unknown or Uncharacterized, BG: I-E+I-P+I-X+UK.

### DECRES gives higher sensitivity and precision on FANTOM annotated regions compared with ChromHMM and ChromHMM-Segway Combined methods

To assess the relative utility of our supervised deep method for CRR prediction, we compared it with the unsupervised ChromHMM and ChromHMM-Segway Combined methods [8, 12] using FANTOM annotations on five available cell types as reference (Fig. 2). It is intuitive that supervised approaches are preferred when labelled training data is sufficient. Furthermore, both unsupervised methods were developed prior to public release of the FANTOM5 data and are therefore at a disadvantage. However, these annotations are widely used by the community and hence the relative performance of DECRES to the standard is of interest. Overall, we observe that DECRES outperforms ChromHMM and Combined methods which in turn deliver similar performance. These unsupervised methods consistently have lower sensitivities for active enhancer detection and lower precision for active promoter detection. Using ChromHMM, the active enhancer sensitivity ranges from 16.9% to 48.4% (numbers are consistent with the test on ENCODE predicted enhancers reported in [14]), while our deep model ranges from 65.9% (K562) to 85.9% (GM12878). Moreover, ChromHMM achieves a maximum precision of 64.8% for active promoter prediction, while the minimum for DECRES is of 81.5%.

**Figure 2:**
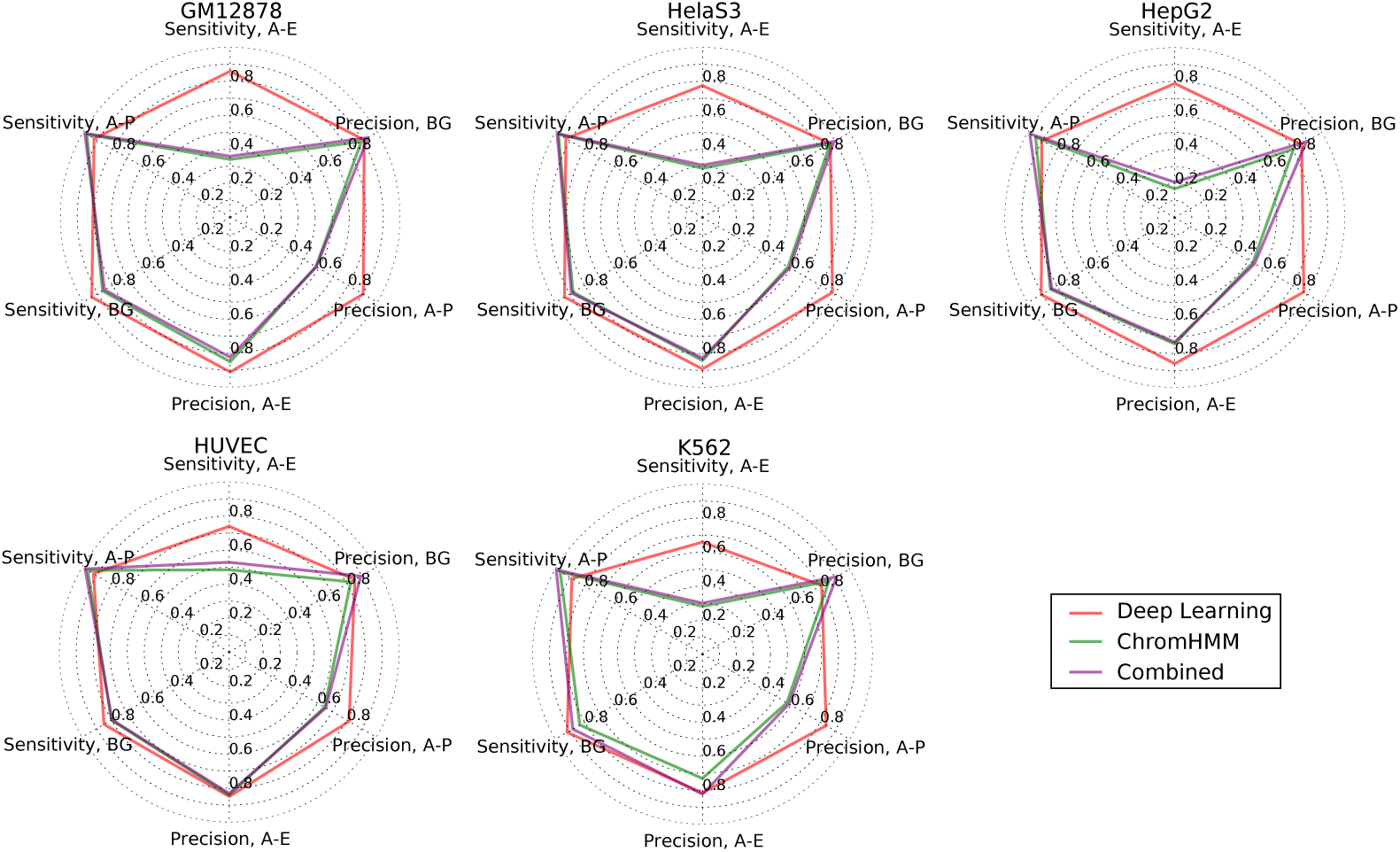
Comparison of the supervised method (Deep Learning) and unsupervised methods (ChromHMM and Combined) on five FANTOM annotated test sets. The ENCODE segmentations were downloaded from http://hgdownload.cse.ucsc.edu/goldenPath/hg19/encodeDCC/wgEncodeAwgSegmentation. We relabelled the annotations of ChromHMM and Combined. For ChromHMM segmentations, the Tss, TssF, and PromF classes were merged to A-P; the Enh, EnhF, EnhW, EnhWF classes were merged to A-E; and the rest were denoted by BG. When processing the Combined annotations, TSS and PF were relabelled to A-P; E and WE were relabelled to A-E; and the rest to BG.

### Evaluation of DECRES performance with independent experimental data

As DECRES is trained on FANTOM regions, we sought two independent collections of laboratory validated enhancers for assessing performance. A collection of predicted enhancers and negative regions (as reported by the Combined ChromHMM and Seqway segmentation method) was previously tested using CRE-seq [14]. In that study, only 33% of predicted regulatory regions were classified as positive in the experiment, compared to 7% for the negative set. Using DECRES trained on all available active regulatory regions of K562 cells, we therefore validated our method on 388 regions showing active enhancer activity in K562 as validated by CRE-seq compared to the 300 control regions (Supplementary Table 3). Highly consistent with the results above, 65.5% (254/388) of the experimentally validated regions were successfully predicted as A-E; the remaining 134 regions were predicted as background (none were classified as promoters). For the 812 tested predictions that were inactive in the Cre-seq experiment, DECRES classified 46.6% (378/812) as positive. For the 300 control regions, DECRES predicted all to be negative (including the 21 that were active in the CRE-seq experiment). Given that the sequences predicted by the Combined method display characteristics that suggestive regulatory activity, the performance of DECRES is satisfactory. We wonder is there any differences between the experimentally supported Combined predictions and the experimentally not-supported Combined predictions. We drew the histogram of DECRES membership scores of 254 and 433 experimentally positive and negative Combined enhancers that were predicted as A-Es by DECRES (Supplementary Fig. 2). It shows that the former set has more certainty (*p* = 0.014, Mann-Whitney rank test).

A collection of 799 HepG2 or K562-specific enhancers defined by motifs of five activators and evolutionary conversation were tested using a massively parallel reporter assay (MPRA) in [33]. In that study, 41% of these enhancers were significantly expressed (*p* = 0.05, Mann-Whitney rank test). We used DECRES to predict the classes of the MPRA positive and MPRA negative enhancers. Our result in Supplementary Table 3 shows that 96.6% (200/207) and 98.5% (123/125) of the MPRA positive enhancers were respectively predicted to be A-Es by DECRES for HepG2 and K562 cells, while 81.7% (223/273) and 92.3% (179/194) of the MPRA negative enhancers were still predicted as A-Es for HepG2 and K562, respectively, but with different (*p* = 6.6E - 7 and *p* = 3.4E - 6 for HepG2 and K562 respectively, Mann-Whitney rank test) distributions of DECRES scores (Supplementary Fig. 2). It reflects an advantage of DRCRES: the class membership scores in DECRES allow user select predictions with levels of confidences. Considering the fact that the negative can include a certain amount of false negatives using *p* = 0.05 as threshold, the DECRES performance on the negative set should not be explained with low specificity.

### Including sequence properties beyond CpG islands does not improve performance for CRR prediction but can inform analysis in the absence of laboratory feature data

Recent studies confirmed that sequence properties can be important for the recognition of promoters and enhancers [3, 5, 24]. It is recognized that the inclusion of CpG islands for promoter analysis improves prediction, primarily for promoters associated to housekeeping genes [34]. For classification, we sought to identify which (if any) additional sequence features contribute to the capacity to distinguish between active promoters and enhancers. We trained the model with 351 sequence features (originally used in [24]) in multiple scenarios. Results are displayed in Figure 3. First, a deep method restricted to sequence features (Fig. 3A) delivered accuracies from 78.11% to 89.28%, confirming that sequence attributes are indeed informative. Second, sequence features have a limited utility for distinguishing between active and inactive states of enhancers (Fig. 3B) and promoters (Fig. 3C), which is logical. For example, on K562, the 10-fold cross-validation accuracy using the sequence properties is only of 68.06%, while the accuracy using the NGS features is as high as 93.85%. Using sequence features in the absence of experimental features has a lower performance across all eight cell types (Fig. 3F). Finally, better results were not achieved by combining experimental and sequence features. For instance taking the 10-fold cross validation of A-E versus A-P versus BG on HelaS3, mean accuracies of 88% and 87.89% were obtained using the experimental features and all features, respectively.

**Figure 3:**
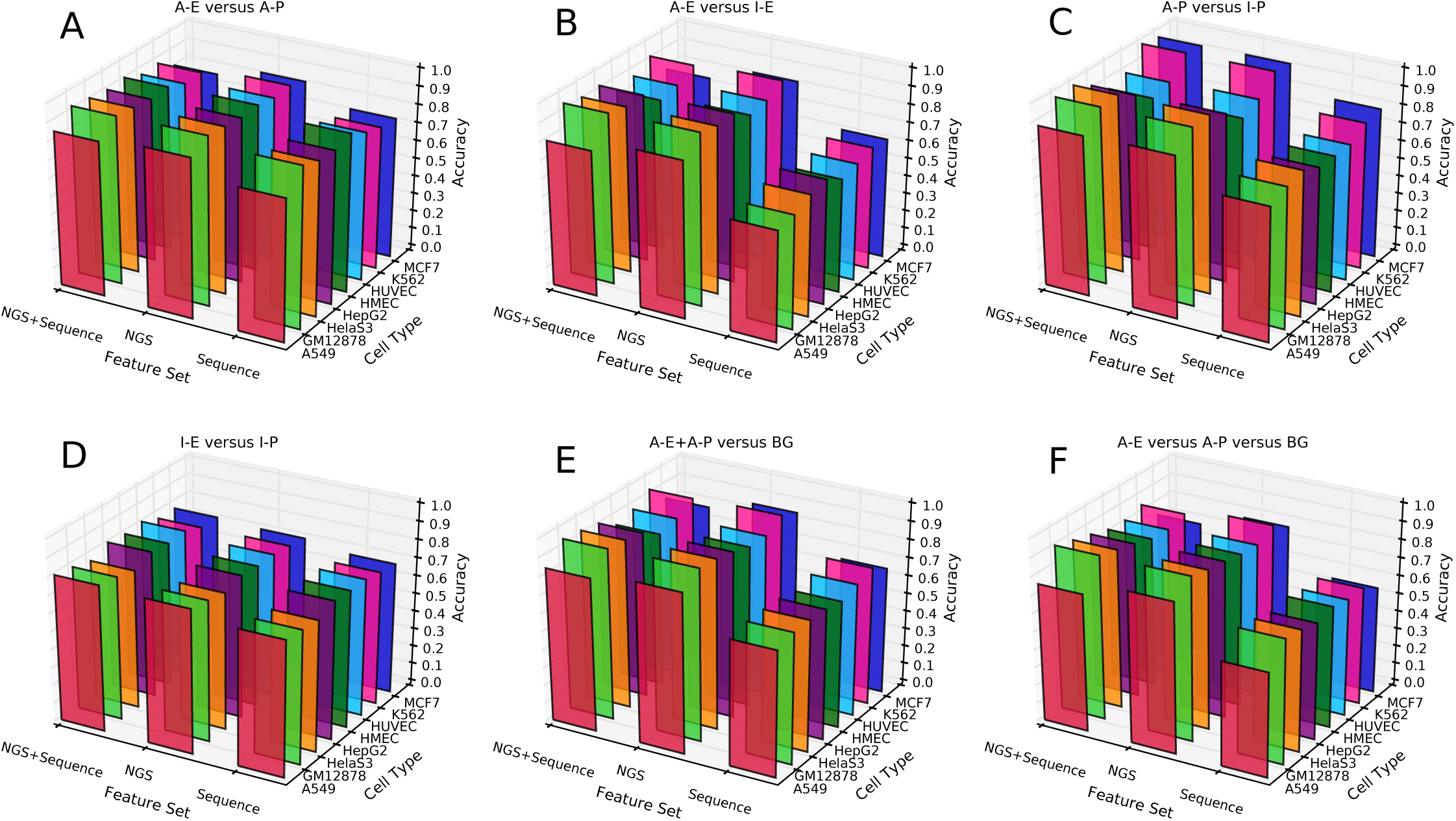
Comparing the 10-fold cross-validation performances on our labelled regions using different feature sets. NGS means our next generation sequencing feature set. Sequence means the set of 351 sequence properties used in [24]. NGS+Sequence means the combination of these two sets.

### A few key features are sufficient

As experimental data can be time consuming and expensive to produce, we sought to determine the minimal set of features most informative for CRR prediction. We used deep feature selection (DFS) models [25] for two-class (A-E+A-P (or CRR) versus BG) and three-class (A-E versus A-P versus BG) classifications on four cell types (GM12878, HelaS3, HepG2, and K562) which have 73-136 features available.

Figures 4A and B show the changes of test accuracies as the numbers of selected features increase for the two-class and three-class classifications, respectively. In both cases, test accuracies increase dramatically for the initial features, then performance stabilizes. A few key features are sufficient for a good prediction accuracy. To define an optimal number of features needed, we fit the curves in Figures 4 and selected the intersection point for a line with slope of 0.25 (see Methods). Fewer features are needed for two-class CRR prediction (4 features) compared to three-class models intended to distinguish between A-E, A-P and background (8 features).

**Figure 4:**
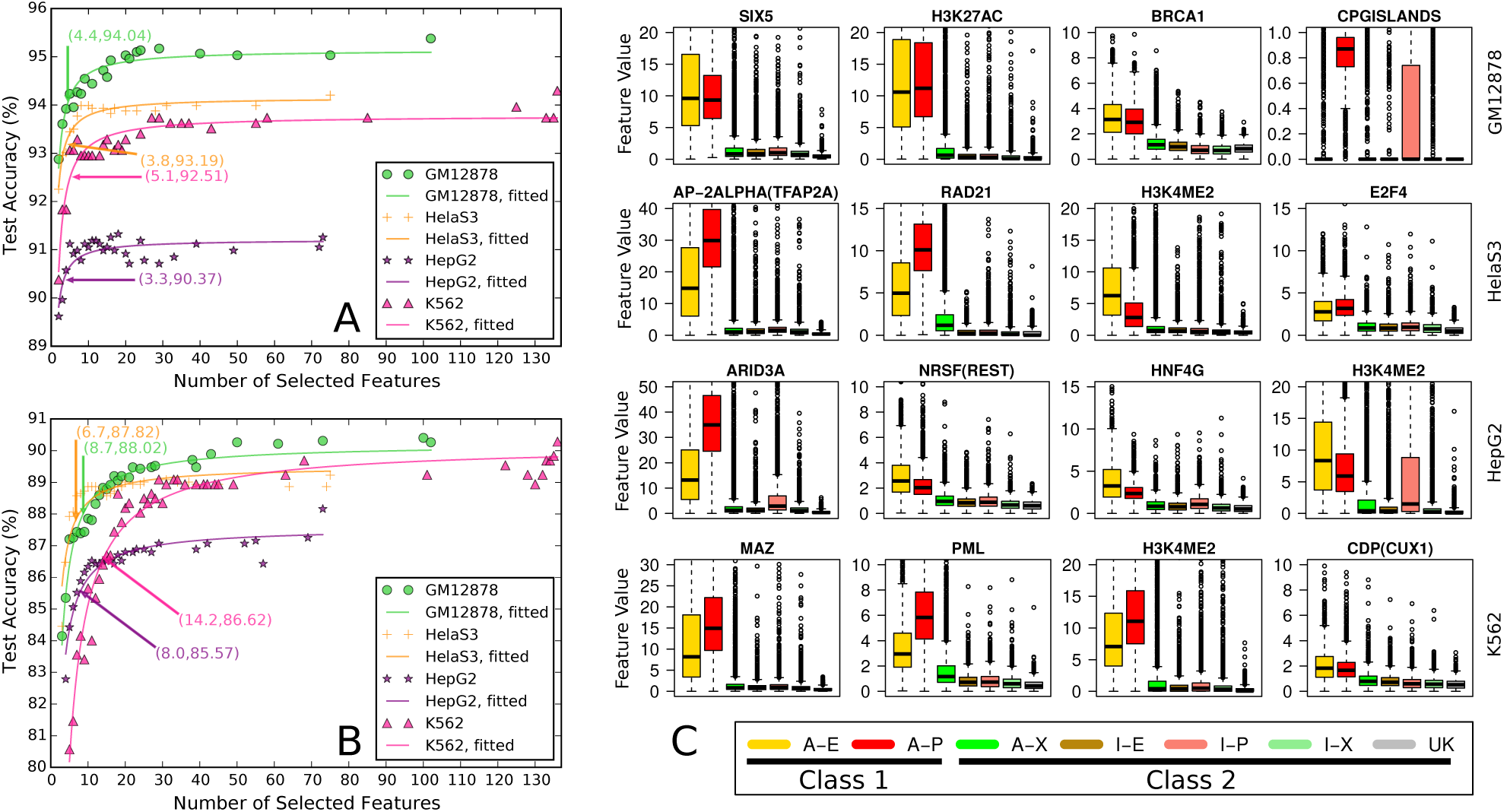
Feature analysis. (A) Accuracy versus the number of features incorporated into the model for 2-class prediction (distinguishing active CRRs (A-E + A-P) from BG (background: A-X, I-E, I-P, I-X and UK). The annotated points indicate where a line with slope 0.25 intersects a fitted curve). (B) Accuracy versus the number of features for a 3-class prediction (distinguishing A-E, A-P and BG). Points as described for (A). (C) For the top 4 features of the 2-class models generated for four well-characterized cell lines, box-plots depict the range of observed feature values (log2 scale) for 7 sequence classes.

Figure 4C shows box plots of the top four selected features for two-class predictions. Using these features, accuracies of 93.92%, 93.01%, 90.57%, and 91.83% were obtained for GM12878, HelaS3, HepG2, and K562, respectively. For most features, the ranges of values are elevated in A-E and A-P relative to the background categories. The majority of the selected features (11 out of 16) are TF binding ChIP-seq data, but each set of four includes a histone modification. The box plots of H3K4me2 (included in 3 out of 4 models) and H3K27ac, indicate that both histone marks distinguish A-E and A-P from background. From the literature, H3K4me2 marks are prevalent at TF bound regions [35], and H3K27ac is enriched at the flanking regions of active CRRs [2].

The distributions of the top eight features for three-class predictions (A-E, A-P, and BG) are given in Supplementary Figure 3. Using the top eight features for each cell, accuracies of 87.39%, 88.65%, 85.88%, and 84.16% were achieved on GM12878, HelaS3, HepG2, and K562, respectively. CpG islands, H3K4me2, and H3K27me3 were commonly selected features for the three-class models, in agreement with existing knowledge that CpG islands mark promoter regions, and inactive regions are enriched for H3K27me3 [36, 2]. Moreover, the well-known enhancer marker H3K4me1 [2] was selected for cell lines GM12878 and HelaS3. H3K4me3 was enriched in both A-E and A-P regions, but had a higher signal level on A-Ps than A-Es, consistent with [3]. The box plots of all features are provided in Supplementary Figures 4-11.

### The majority of DECRES’s genome-wide predictions are supported by other methods

We trained 2- and 3-class multilayer perceptron (MLP) models (see Methods) using all reference (labelled) data for training, in order to predict CRRs across the entire genome for six cell types (A549 and MCF7 were excluded). The 2-class model identified 227,332 CRRs (adjacent regions were merged), which occupy 4.8% of the genome (Supplementary Table 4). A total of 9,153 CRRs were ubiquitously predicted across all six cell types. For the 3-class prediction, we obtained 301,650 A-E regions (6.8% of the genome) and 26,555 A-P regions (0.6% of the genome) together with 11,886 ubiquitous A-Es and ubiquitous 3,678 A-Ps. The genome-wide predictions for all six cell types are available in Supplementary Data 1.

Next, we examined the overlap of our predicted CRRs with the Combined [12] and dReg [21] predictions on GM12878, HelaS3, and K562 (Fig. 5A, B, and C). The majority of CRRs predicted by DECRES overlap with the results from either Combined or dReg, specifically 86.13%, 76.13%, and 83.63% for GM12878, HelaS3, and K562, respectively (Fig. 5A, B, and C). A subset (13.87% on GM12878, 23.87% on HelaS3, and 16.37% on K562) of DECRES predictions do not overlap with predictions from the other two tools. Notably, a large portion of the Combined predictions (56.78% on HelaS3, 55.99% on GM12878, and 36.36% on K562) do not overlap with those from the supervised methods, which is consistent with its low observed validation rate [14]. DECRES predictions (see Fig. 5D for an example) have two distinguishing characteristics: a finer resolution for both A-P and A-E regions, and the A-P regions tend to be wrapped by A-E regions. Indeed, Fig. 5E shows that 51.73% - 70.75% of A-Ps are surrounded by A-Es, with the exception of K562 where 95.75% of A-Ps are wrapped by A-Es. It is unclear why K562 promoters display such strong properties.

**Figure 5:**
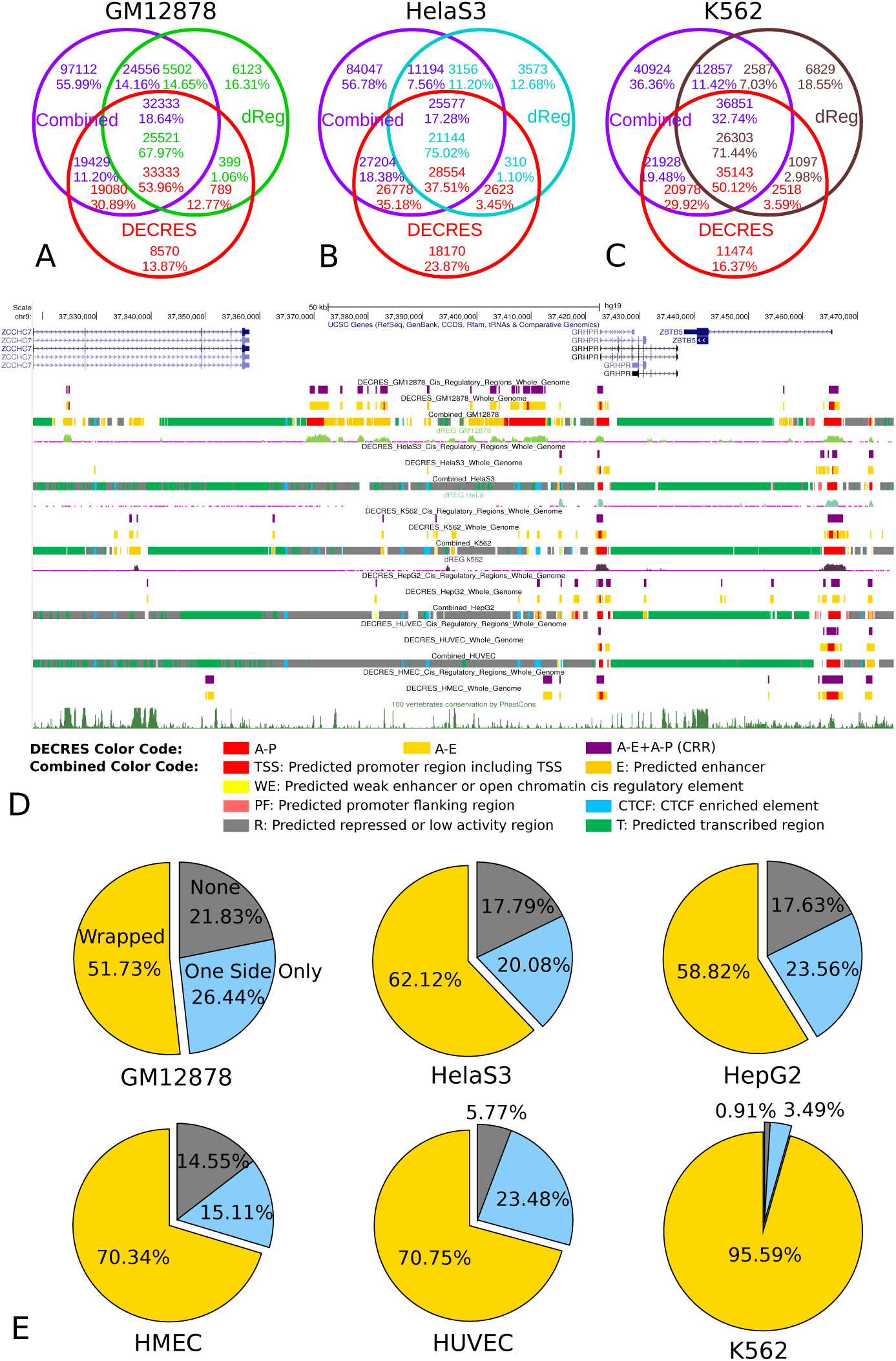
Comparing the genome-wide predictions with Combined and dReg. (A,B,C) The agreements of the DECRES CRRs with the Combined and dReg CRRs. The TSS, PF, E, and WE segmentations from Combined were relabelled to CRRs. The active transcriptional regulatory elements (TREs) predicted by dReg were renamed to CRRs. (D) An example of predicted regulatory regions in the UCSC Genome Browser. (E) Proportions of active promoters wrapped by active enhancers. Wrapped: a promoter prediction is surrounded by enhancer predictions from both sides within a 1 kbp extension respectively. One Side Only: enhancer predictions appear only at either the left or right-side (but not both sides) of a promoter prediction (within a 1 kbp extension). None: no enhancer prediction falls in the 1 extensions of a promoter prediction.

We investigated how many among our genome-wide predictions are supported by the VISTA enhancer set [37]. Despite that the majority of the VISTA enhancers are extremely conserved for development, we still find that 37.1% (850/2,293) of experimentally confirmed and unconfirmed VISTA enhancers overlap with the predicted A-Es, while merely 4.8% (110/2,293) of these VISTA enhancers overlap with the predicted A-Ps. Results for experimentally confirmed VISTA enhancers are similar (482/1,196=40.30% and 60/1,196=5.02% overlap A-Es and A-Ps, respectively), which suggests that our predicted active enhancers have real enhancer functions. A proportion of the VISTA enhancers not overlapping our predictions could be active specifically in other cell types than our focus cell lines.

### Extending the FANTOM enhancer atlas

Due to the limited depth of CAGE signals for eRNAs, a portion of active (or transcribed) enhancers will not have been detected in the original compilation of the enhancer atlas. Hence, we sought to identify additional partially supported enhancers for which eRNA signals were below the original atlas threshold settings [4]. In the previous work, a total of 200,171 bidirectionally transcribed (BDT) loci were detected across the human genome, using CAGE tags of 808 cell types and tissues. After excluding BDT loci within exons, a partially supported set of 102,021 BDT regions remained, of which 43,011 balanced loci (similar eRNA levels on both sides) constitute the FANTOM enhancer atlas [4]. In order to investigate whether more active enhancer candidates can be detected for each of the six cell types, we trained a MLP on its active atlas regions, and predicted classes for all 102,021 BDT sites. Among the 102,021 BDT loci, most were classified as negative regions in a given cell (Supplementary Table 5), while on average 13,316 were predicted as A-Es and only 834 were predicted as A-Ps per cell type. A substantial number (6,535 on average) of inactive enhancers in the original enhancer atlas were predicted as active by our model (see Supplementary Table 6), consistent with the assumption that BDT data is incomplete for any given sample. On average 5,514 BDT loci excluded by the original atlas, were predicted as A-Es per cell type. Over the six analyzed cell types, a total of 38,601 BDT loci were predicted as A-Es (Supplementary Data 2), of which 16,988 represent an expansion of the original FANTOM enhancer atlas. Note that 21,398 out of 43,011 enhancers from the original FANTOM enhancer atlas are not predicted as active in the six cells analyzed here, but these regions may be active in the other 802 cells for which there are inadequate features to analyze.

### Functional and motif enrichment analysis

We performed functional enrichment analysis on the genome-wide predicted A-Es and A-Ps using GREAT [38]. For GM12878 cells, 79% of predicted enhancer regions are more than 5 kilobase pairs (kbps) away from gene TSSs (Fig. 6A), while 47% of predicted promoters are less than 5 kbps to the annotated gene TSSs (Fig. 6B). Similar statistics were obtained for the remaining five cell types. Annotation analyses of the GM12878-specific CRRs shows that proximal genes are associated to: immune response from gene ontology (GO) annotations (Fig. 6C); B cell signalling pathways from MSigDB Pathway annotations (Fig. 6D); and leukemia from disease ontology annotations (Fig. 6E). Results are consistent with the lymphoblastoid lineage of the cells. Next, we performed functional enrichment analysis on the BDT-supported predicted enhancers not previously reported in the FANTOM enhancer atlas (NiA, “not in atlas”). Results are fully consistent with the above analysis (Supplementary Fig. 12).

**Figure 6:**
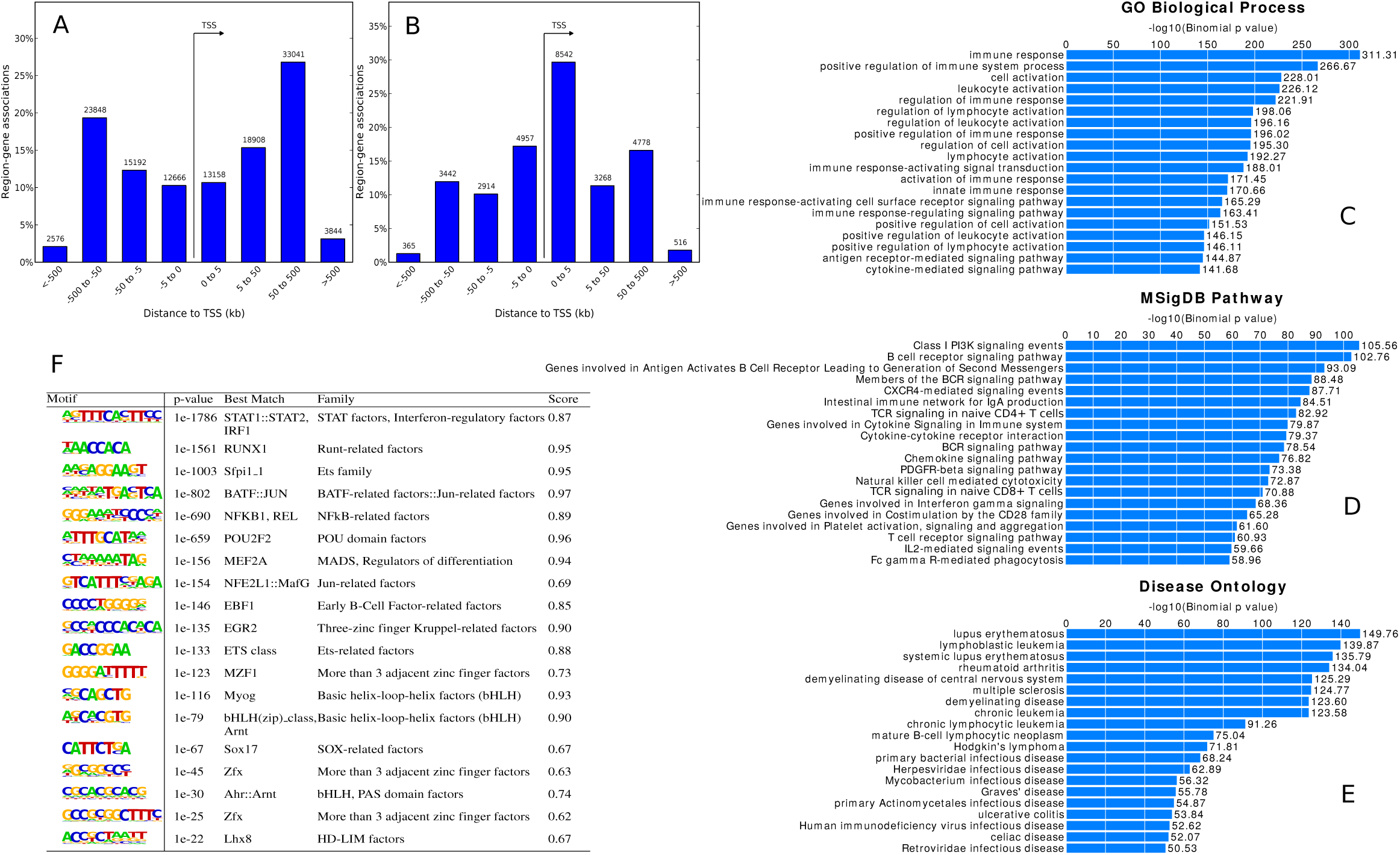
Functional and motif analysis of the genome-wide predictions on cell line GM12878. (A) The distance from the predicted A-Es to gene TSSs. (B) The distance from the predicted A-Ps to gene TSSs. (C,D,E) Top 20 enriched biological processes, pathways, and diseases, respectively, in the predicted cell-specific CRRs. (F) Enriched *de novo* motifs in the predicted cell specific CRRs. Column 4: the families of best-matched TFs. Column 5: the best match scores.

We further carried out motif enrichment analysis on the predicted cell-specific CRRs and NiA enhancers using HOMER [39]. The predicted regions are enriched for motifs similar to JASPAR binding profiles [40] (Fig. 6F and Supplementary Fig.s 12-22) both associated to TFs maintaining general cell processes and TFs with selective roles in cell-related functions. For instance, motifs for Jun-, Fos-, and Ets-related factors were enriched in regions from all six cell types (Fig. 6F, Supplementary Fig.s 12-22). These TFs regulate general cellular progresses such as differentiation, proliferation, or apoptosis [41, 42]. Cell-appropriate TF enrichments were observed for each cell (summarized in Supplementary Table 7). For example, RUNX1 and other Runt-related factors, which play crucial roles in haematopoiesis, are observed in GM12878 (Fig. 6F and Supplementary Fig. 12) [43]. C/EBP-related factors that regulate genes involved in immune and inflammatory responses are expressed in cervix (Supplementary Fig.s 13 and 14) [44]. HNF1A, HNF1B, FOXA1, FOXA2, HNF4A, and HNF4G factors regulate liver-specific genes (Fig.s 13 and 14) [45, 46]. NFY factors cooperate with GATA1 to mediate erythroid-specific transcription in K562 (Supplementary Fig.s 21 and 22) [47].

## Discussion and conclusion

We show that using FANTOM data for training, supervised deep learning methods are able to accurately predict active enhancers and promoters across the human genome. We demonstrate that cell-specific data outperforms universal data (e.g. sequence), and highlight key experimental features that tend to be incorporated into predictive models when available. We explore the relative performance of 2- and 3-class models to group or separate enhancers and promoters. Finally, we deliver a comprehensive collection of annotations, that label 6.8% of the genome as enhancers and 0.6% as promoters in one or more of six well-characterized cells.

Accurate annotation of regulatory regions across the human genome is essential for genome interpretation. With genome sequencing transitioning to a standard clinical test in the next few years, the ability to move beyond the analysis of protein-coding alterations has the potential to greatly enhance our capacity to explain observed genetic disorders. By demonstrating the suitability of supervised deep learning methods to provide such labels, we now enter into a new stage of genome annotation. In the next few years, we can anticipate that laboratories working with specific cell types will compile genome-scale feature data and subsets of eRNA-supported regulatory sequences, allowing the broad identification of reliable regulatory regions. As raised in the introduction and demonstrated in this report, deep learning methods are well suited for this challenge.

Our study brings together a large collection of high-throughput data from global projects to allow for supervised annotation. One key challenge in such analysis is the depth of validation performed. In this report, validation is assessed using existing collections of reliable enhancers, including CAGE [4], and laboratory validated sets from CRE-seq [14], and transgenic mouse assays [37]), showing that the supervised approach nears 85.9% sensitivity. The capacity to use supervised approaches is important, as unsupervised approaches are limited. While we compare to multiple laboratory validated sets retrospectively, a prospective assessment would have broad value. Given the cost of such prospective testing, we propose that due to the arrival of sufficient data for supervised approaches on a large-scale, it is time to conduct a global prospective assessment, such as enabled within the DREAM Challenge program (http://dreamchallenges.org). Such a test for annotation of *cis*-regulatory regions in the human genome would inspire the machine learning community to push the performance limit of supervised CRR-prediction methods.

Enhancers and promoters have both common and distinct characteristics. In our cross-validations, we show that A-E and A-P are highly separable (Fig. 1A), while better mean accuracy can be obtained if A-E and A-P are merged together rather than being treated separately (Fig. 1E and F). Both continuous (merging enhancers and promoters together) and distinct models (treating enhancers and promoters separately) have limitations. While a continuous model may overlook functional differences, a distinct model may overemphasize such differences. A potentially better prediction model might require two hierarchical steps. It could first distinguish CRRs from the background genome, then assign a continuous score to each candidate region indicating the likelihood of being an enhancer. Further clustering and subtyping may be necessary.

Two other deep learning models might be beneficial to improve annotations of non-coding regions. One method is convolutional neural network, which can take into account the shape information of various features. The other one is bidirectional recurrent neural network, which can consider the information from adjacent context. It can be potentially applied to annotate regulatory domains or complexes where exons, introns, promoters, enhancers, silencers, and insulators form cohorts for specific functionalities.

We anticipate that the collection of regulatory region annotations provided in this study will have broad utility for genome interpretation, and that the demonstration of the sufficiency of training data and the utility of deep learning supervised methods for CRR prediction will move the discussion to a highly applied period of high-quality annotation. Understanding how CRRs interact and how they link to their target genes is the key to decipher the cis-regulatory mechanism. We expect that further development of integrative machine learning methods is crucial to reconstruct such a gene regulatory system.

## Methods

### Data

For the purpose of supervised analysis, we collected feature data from ENCODE [10] along with the transcriptionally active enhancers and promoters from eight matched cell types catalogued by the FANTOM effort [23, 4]. These cell types include A549, GM12878, HelaS3, HepG2, HMEC, HUVEC, K562, and MCF7. For each cell type, we defined seven classes of labelled regions, including A-E, I-E, A-P, I-P, A-X, I-X, and UK. The libraries of enhancers and promoters were downloaded from http://fantom.gsc.riken.jp/5/data. A-Es and I-Es were defined as FANTOM enhancers with TPM>0 (tags per million) and TPM=0, respectively. A-Ps and I-Ps were randomly selected FANTOM promoters with TPM>5 and TPM=0, respectively. A-X and I-X were defined based on exons’ transcription levels measured by RNA-seq (ftp://hgdownload.cse.ucsc.edu/goldenPath/hg19/encodeDCC). An exon with peak-max greater than 200 (equal to 0) was defined as A-X (I-X). The UK regions were sampled from the genome regions excluding all FANTOM CAGE tags, all exons, and DNaseI open regions. The numbers of labelled regions used in this study are listed in Supplementary Table 1. For each cell type, we built a comprehensive feature set, integrating histone modification and TF binding ChIP-seq, DNase-seq, RNA-seq, FAIRE-seq, and ChIA-PET data from the ENCODE project (http://www.broadinstitute.org/~anshul/projects/encode/rawdata/signal/mar2012/pooledReps/bigwig/macs2signal/foldChange and ftp://hgdownload.cse.ucsc.edu/goldenPath/hg19/encodeDCC).

These features characterize the activities of enhancers and promoters in cell-specific aspects. Additionally, CpG islands and phastCons evolutionary conservation scores were included, because it is well recognized that some regulatory regions are highly GC-rich and extremely conserved. For each labelled region, the mean value of feature signals fall within a bin centered at the region was taken as the feature value using bwtool [48].

We tried different bin sizes including 200, 500, 1,000, 2,000, and 4,000 bps. Since 200 bps worked the best in cross-validation tests, we used it throughout our analyses. The numbers of features used for each cell type and the numbers of common features between any pair of them are given in Supplementary Table 2. A combined list of features is provided in Supplementary Data 3. Our labelled data are downloadable from https://github.com/yifeng-li/DECRES.

### Deep learning for classification

Based on the Deep Learning Tutorials (deeplearning.net/tutorial) and Theano (deeplearning.net/software/theano), we implemented a deep learning package named DECRES (DEep learning for identifying Cis-Regulatory ElementS and other applications) which is available at https://github.com/yifeng-li/DECRES. We applied a supervised deep model - feedforward neural network (also known as multilayer perceptrons or MLP) for the detection of regulatory regions. Along the data flow in the model structure, it has three hidden layers with 256, 128, and 64 hidden units, respectively. In order to control model complexity, l2-norm was used as a regularization term with control parameter 0.01. The maximum number of allowed iterations was set to 1,000. The initial learning rate was set to 0.1, and was reduced as the number of iterations increases. Using batch-size 100, stochastic gradient descent was employed to optimize the model parameters. The momentum term with control parameter 0.1 was added to the update rule in order to stabilize the optimization. Hyperbolic tangent activation function was used. It is well-known that the model selection of neural networks is time consuming, we thus manually selected the above control parameter setting by trying a variety of settings, rather than resort to an automatic model searching method though it may further improve the performance. When evaluating the classification performance of various experiments, 10-fold cross-validation was used to split a labelled data set into training and test sets. A training set was further partitioned into a training subset for model learning and a validation subset for early termination. Before predicting regulatory regions in the whole genome, all labelled data of a cell type were used to train the network.

### Feature selection

Our newly devised deep feature selection (DFS) model [25] was used to select subsets of discriminative features. Addressing the limitations of sparse linear models for feature selection, DFS is able to model the non-linearity of the features and select a single subset of features for multi-class data. The main idea of DFS is to add a one-to-one linear layer (named feature-selection layer) to the above described feedforward neural network. For the *i*-th input feature *x*_*i*_, the output of the feature-selection layer becomes *w*_*j*_*x*_*j*_. Thus, the parameter of this layer is a vector *w*. By shrinking *w*, some of its elements turn to zeros, such that the corresponding features do not contribute to the classification at all. The upper hidden layers of the model have the capability of modelling the non-linear interactions in the data. The feature selection layer allows to select a single subset of features for multi-class problem. Aiming at trading off the size of feature subset and accuracy, we designed a method based on curving fitting that was applied in Fig. 4. Denoting a size of feature subset and corresponding test accuracy by *x* and *y* respectively, we first fit function 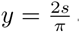 arctan(*kx*) where *k* and *s* are scale parameters. Once done, a point can be chosen on the curve given a proper tangent value (say *t*) using 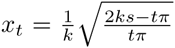 and 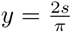 arctan(*kx*_*t*_). Since the values come from different scales on *x* and *y* axes, we qualitatively used *t* = 0.25. The DFS model and described feature-subset-selection method were included in the DECRES package.

## Acknowledgments

We thank our colleagues in the Wasserman Laboratory for discussions and feedback improving the quality of this paper. We thank Dr. Anshul Kundaje (Stanford) for making the processed ChIP-seq fold-change data and script available. The Wasserman laboratory is supported by the Genome Canada/Genome BC: 174DE (ABC4DE project), Canadian Institutes of Health Research (CIHR): MOP-82875, the Natural Sciences and Engineering Research Council of Canada (NSERC): RGPIN355532-10, and the National Institutes of Health (USA) grant: 1R01GM084875. The computer systems of the Gene Regulation Bioinformatics Laboratory were funded by the Canada Foundation for Innovation and the BC Knowledge Development Fund. WWW was supported by scholar awards from CIHR and the Michael Smith Foundation for Health Research (MSFHR). YL was supported by NSERC Postdoctoral Fellowship: PDF-471767-2015.

## Author contributions

Y.L. contributed the implementation, computational experiments, and the initial draft. W.S. compiled experimental enhancer sets and provided feedback on the project and manuscript. W.W.W. supervised the whole project and contributed to the results analysis and manuscript writing.

## Competing interests

The authors declare that they have no competing financial interests.

## References

[1]. Pennacchio L., Bickmore W., Dean A., Nobrega, M. & Bejerano, G. Enhancers: Five essential questions. Nature Review Genetics 14, 288–295 (2013).

[2]. Shlyueva D., Stampfel, G. & Stark, A. Transcriptional enhancers: From properties to genome-wide predictions. Nature Review Genetics 15, 272–286 (2014).

[3]. Andersson R., Sandelin, A. & Danko, C. A unified architecture of transcriptional regulatory elements. Trends in Genetics 31, 426–433 (2015).

[4]. Andersson, R. et al. An atlas of active enhancers across human cell types and tissues. Nature 507, 455–461 (2014).

[5]. Core, L. et al. Analysis of nascent RNA identifies a unified architecture of initiation regions at mammalian promoters and enhancers. Nature Genetics 46, 1311–1320 (2014).

[6]. Wasserman, W. & Sandelin, A. Applied bioinformatics for the identification of regulatory elements. Nature Review Genetics 5, 276–287 (2004).

[7]. Li Y., Chen C., Kaye, A. & Wasserman, W. The identification of cis-regulatory elements: A review from a machine learning perspective. BioSystems 138, 6–17 (2015).

[8]. Ernst, J. & Kellis, M. ChromHMM: Automating chromatin-state discovery and characterization. Nature Methods 9, 215–216 (2012).

[9]. Hoffman, M. et al. Unsupervised pattern discovery in human chromatin structure through genomic segmentation. Nature Methods 9, 473–476 (2012).

[10]. The ENCODE Project Consortium. An integrated encyclopedia of DNA elements in the human genome. Nature 489, 57–74 (2012).

[11]. Johnson D., Mortazavi A., Myers, R. & Wold, B. Genome-wide mapping of in vivo protein-DNA interactions. Science 316, 447–455 (2007).

[12]. Hoffman, M. et al. Integrative annotation of chromatin elements from ENCODE data. Nucleic Acids Research 41, 827–841 (2013).

[13]. Kwasnieski, J. et al. Complex effects of nucleotide variants in a mammalian cis-regulatory element. Proceedings of the National Academy of Sciences 109, 19498–19503 (2012).

[14]. Kwasnieski J., Fiore C., Chaudhari, H. & Cohen, B. High-throughput functional testing of ENCODE segmentation predictions. Genome Research 24, 1595–1602 (2014).

[15]. Yip, K. et al. Classification of human genomic regions based on experimentally determined binding sites of more than 100 transcription-related factors. Genome Biology 13, R48 (2012).

[16]. Rajagopal, N. et al. RFECS: A random-forest based algorithm for enhancer identification from chromatin state. PLOS Computational Biology 9, e1002968 (2013).

[17]. Chen C., Morris, Q. & Mitchell, J. Enhancer identification in mouse embryonic stem cell using integrative modeling of chromatin and genomic features. BMC Genomics 13, 152 (2012).

[18]. Arnold, C. et al. Genome-wise quantitative enhancer activity maps identified by STARR-seq. Science 339, 1074–1077 (2013).

[19]. Yanez-Cuna, J. et al. Dissection of thousands of cell type-specific enhancers identifies dinucleotide repeat motifs as general enhancer features. Genome Research 24, 1147–1156 (2014).

[20]. Core L., Waterfall, J. & Lis, J. Nascent RNA sequencing reveals widespread pausing and divergent initiation at human promoters. Science 322, 1845–1848 (2008).

[21]. Danko, C. et al. Identification of active transcriptional regulatory elements from GRO-seq data. Nature Methods 12, 433–438 (2015).

[22]. Kodzius R., Kojima M., Nishiyori, H. et al. CAGE: Cap analysis of gene expression. Nature Methods 3, 211–222 (2006).

[23]. The FANTOM Consortium, The RIKEN PMI & CLST (DGT). A promoter-level mammalian expression atlas. Nature 507, 462–470 (2014).

[24]. Kleftogiannis D., Kalnis, P. & Bajic, V. DEEP: A general compuational framework for predicting enhancers. Nucleic Acids Research 43, e6 (2015).

[25]. Li Y., Chen, C. & Wasserman, W. Deep feature selection: Theory and application to identify enhancers and promoters. Journal of Computational Biology in press, DOI:10.1089/cmb.2015.0189 (2015).

[26]. Hinton G., Osindero, S. & Teh, Y. A fast learning algorithm for deep belief nets. Neural Computation 18, 1527–1554 (2006).

[27]. Hinton, G. & Salakhutdinov, R. Reducing the dimensionality of data with neural networks. Science 313, 504–507 (2006).

[28]. Bengio Y., Courville, A. & Vincent, P. Representation learning: A review and new perspectives. IEEE Transactions on Pattern Analysis and Machine Intelligence 35, 1798–1828 (2013).

[29]. LeCun Y., Bengio, Y. & Hinton, G. Deep learning. Nature 521, 436–444 (2015).

[30]. Xiong, H. et al. The human splicing code reveals new insights into the genetic determinants of disease. Science 347, 1254806 (2015).

[31]. Alipanhi B., Delong A., Weirauch, M. & Frey, B. Predicting the sequence specificities of DNA-and RNA-binding proteins by deep learning. Nature Biotechnology 33, 831–838 (2015).

[32]. Spencer M., Eickholt, J. & Cheng, J. A deep learning network approach to ab initio protein secondary structure prediction. IEEE/ACM Transactions Computational Biology and Bioinformatics 12, 103–112 (2015).

[33]. Kheradpour, P. et al. Systematic dissection of regulatory motifs in 2000 predicted human enhancers using a massively parallel reporter assay. Genome Research 23, 800–811 (2013).

[34]. Deaton, A. & Bird, A. CpG islands and the regulation of transcription. Genes & Development 25, 1010–1022 (2011).

[35]. Wang Y., Li, X. & Hua, H. H3K4me2 reliably defines transcription factor binding regions in different cells. Genomics 103, 222–228 (2014).

[36]. Zhou V., Goren, A. & Bernstein, B. Charting histone modifications and the functional organization of mammalian genomes. Nature Review Genetics 12, 7–18 (2011).

[37]. Visel A., Minovitsky S., Dubchak, I. & Pennacchio, L. VISTA Enhancer Browser – a database of tissue-specific human enhancers. Nucleic Acids Research 35, D88–D92 (2007).

[38]. McLean, C. et al. GREAT improves functional interpretation of cis-regulatory regions. Nature Biotechnology 28, 495–501 (2010).

[39]. Heinz S., Benner C., Spann N., Bertolino, E. et al. Simple combinations of lineage-determining transcription factors prime cis-regulatory elements required for macrophage and B cell identities. Molecular Cell 38, 576–589 (2010).

[40]. Mathelier, A. et al. JASPAR 2016: A major expansion and update of the open-access database of transcription factor binding profiles. Nucleic Acids Research 44, D110–D115 (2016).

[41]. Ameyar M., Wisniewska, M. & Weitzman, J. A role for AP-1 in apoptosis: The case for and against. Biochimie 85, 747–752 (2003).

[42]. Sharrocks, A. The ETS-domain transcription factor family. Nature Reviews Molecular Cell Biology 2, 827–837 (2001).

[43]. Okuda T., Nishimura M., Nakao, M. & Fujita, Y. RUNX1/AML1: A central player in hematopoiesis. International Journal of Hematology 74, 252–257 (2001).

[44]. Arnett B., Soisson P., Ducatman, B. & Zhang, P. Expression of CAAT enhancer binding protein beta (C/EBP beta) in cervix and endometrium. Molecular Cancer 2, 21 (2003).

[45]. Costa R., Kalinichenko V., Holterman, A. & Wang, X. Transcription factors in liver development, differentiation, and regeneration. Hepatology 38, 1331–1347 (2003).

[46]. Wang Z., Bishop, E. & Burke, P. Expression profile analysis of the inflammatory response regulated by hepatocyte nuclear factor 4α. BMC Genomics 12, 128 (2011).

[47]. Fleming, J. et al. NF-Y coassociates with FOS at promoters, enhancers, repetitive elements, and inactive chromatin regions, and is stereo-positioned with growth-controlling transcription factors. Genome Research 23, 1195–1209 (2013).

[48]. Pohl, A. & Beato, M. bwtool: A tool for bigWig files bioinformatics. Bioinformatics 30, 1618–1619 (2014).

